# Unique Genotypic Features of HIV-1 C Gp41 Membrane Proximal External Region Variants during Pregnancy Relate to Mother-to-child Transmission via Breastfeeding

**DOI:** 10.1101/2020.09.07.286278

**Authors:** Li Yin, Kai-Fen Chang, Kyle J. Nakamura, Louise Kuhn, Grace M. Aldrovandi, Maureen M. Goodenow

## Abstract

Mother-to-child transmission (MTCT) through breastfeeding remains a major source of pediatric HIV-1 infection worldwide. To characterize plasma HIV-1 subtype C populations from infected mothers during pregnancy that related to subsequent breast milk transmission, an exploratory study was designed to apply next generation sequencing and a custom bioinformatics pipeline for HIV-1 gp41 extending from heptad repeat region 2 (HR2) through the membrane proximal external region (MPER) and the membrane spanning domain (MSD). Viral populations during pregnancy from women who transmitted by breastfeeding, compared to those who did not, displayed greater biodiversity, more frequent amino acid polymorphisms, lower hydropathy index and greater positive charge. Viral characteristics were restricted to MPER, failed to extend into flanking HR2 or MSD regions, and were unrelated to predicted neutralization resistance. Findings provide novel parameters to evaluate an association between maternal MPER variants present during gestation and lactogenesis with subsequent transmission outcomes by breastfeeding.

**IMPORTANCE:** HIV-1 transmission through breastfeeding accounts for 39% of MTCT and continues as a major route of pediatric infection in developing countries where access to interventions for interrupting transmission is limited. Identifying women who are likely to transmit during breastfeeding would focus therapies during the breastfeeding period to reduce MTCT. Findings from our pilot study identify novel characteristics of gestational viral MPER quasispecies related to transmission outcomes and raise the possibility for predicting MTCT by breastfeeding based on identifying mothers with high-risk viral populations.

## INTRODUCTION

Mother-to-child HIV-1 transmission (MTCT) can occur during pregnancy, delivery (perinatally) or by breastfeeding and contributes substantially to global morbidity and mortality for children under-5 years of age. Rates of perinatal MTCT range from 15% to 45% in the absence of any interventions, but can be reduced to less than 5% with appropriate antiretroviral treatment (1–5). HIV-1 transmission through breastfeeding accounts for 39% of MTCT and continues as a major route of pediatric infection in developing countries (6), when access to interventions for interrupting transmission are limited (7).

Viruses that establish MTCT either perinatally or through breastfeeding display limited diversity, as well as relatively short and under-glycosylated gp120 regions (8–12), similar to gp120 regions among transmitter/founder viruses in general (13–16). The membrane-proximal external region (MPER) of gp41 contains linear epitopes for broadly HIV-1 neutralizing antibodies (bn-HIV-Abs), e.g. 2F5, 4E10, Z13, Z13e1 and 10E8, and is accessible to plasma bn-HIV-Abs (17–21). Elevated maternal antibody titers to HIV-1 envelope (*env*) gp41 and/or gp120 epitopes are directly associated with perinatal MTCT (22–26). Our previous study of HIV-1 MPER sequences from HIV-1 infected mother-baby pairs in the Zambia Exclusive Breastfeeding Study (ZEBS), a clinical trial to prevent MTCT of HIV-1 through breast milk (27–29), suggests that polymorphisms in MPER occur naturally and can confer resistance to broadly neutralizing anti-MPER antibodies (29). Thus, it is plausible to hypothesize that HIV-1 MPER variants in mothers who transmit HIV-1 to their babies by breastfeeding (TM) display a greater extent of genetic polymorphism in MPER compared to those who do not transmit (NTM).

Cross-sectional as well as longitudinal studies of cell-free HIV-1 find persistent mixing and synchronous evolution of viruses between plasma and breast milk in the ZEBS and other cohorts indicating that HIV-1 quasispecies in plasma are representative of virus populations in breast milk (27,30–34), although compartmentalization of cell-associated viruses in breast milk is reported in other studies (30,35). A sophisticated phylogenetic analysis of longitudinal HIV-1 *env* V1-V5 sequences from plasma and breast milk of transmitting mothers suggests that the most common ancestral virus(es) in breast milk originate during the second or third trimester of pregnancy, close to the onset of lactogenesis (27). Consequently, plasma HIV-1 variants during pregnancy might harbor genetic features related to subsequent breast milk transmission.

To examine the relationship between maternal viruses during gestation and subsequent transmission outcomes through breastfeeding, a pilot study of ZEBS maternal plasma subtype C HIV-1 from second or third trimester of pregnancy were evaluated by next generation sequencing (NGS) to provide broad coverage of HIV-1 quasispecies at the population level and sensitive detection of low-frequency variants. A custom bioinformatic pipeline was developed to assess biodiversity, amino acid substitutions within linear epitopes of known bn-HIV-Abs targeting gp41 MPER, and biochemical features (hydropathy and charge) of plasma subtype C HIV-1 gp41 MPER variants, and compared to the adjacent heptad repeat region 2 (HR2) or membrane spanning domain (MSD) among mothers who transmitted or did not transmit HIV-1 through breastfeeding.

## MATERIALS AND METHODS

### Study cohort

A nested, case-control study included a subset of eight women infected by subtype C HIV-1 enrolled in ZEBS (27–29). All subjects were therapy-naïve, except for a single peripartum dose of nevirapine according to the Zambian government guidelines during the enrollment period (2001 - 2004). Written informed consent for participation in the ZEBS study was obtained from all participants. From the larger cohort, our study included plasma samples from four women who transmitted HIV-1 during the early breastfeeding period (TM) (defined by infants who became HIV-1 DNA positive after 42 days following prior negative tests), and four infected women who did not to transmit (NTM) [defined by infants who remained HIV-1 DNA negative through the completion of all breastfeeding for a median (quartile range) of 6.5 (4.0 – 18.8) months)] (Table 1). Maternal plasma samples were collected prospectively during the second/third trimester of pregnancy [median (quartile range): 80 (32 - 164) days before delivery] (Table 1). At the time of sampling, the two groups of women were balanced for median (quartile range) of age [TM, 25.5 (22.5 - 31.5) years vs. NTM, 27.0 (20.3 - 34.5) years] (p = 0.87), CD4 T-cell count [TM, 146 (117 - 187) cells/μl vs. NTM, 202 (132 - 240) cells/μl] (p = 0.27), plasma viral load [TM, log_10_ 5.2 (4.9 - 5.5) HIV-1 RNA copies/ml plasma vs. NTM, log_10_ 5.2 (5.0 - 5.3) HIV-1 RNA copies/ml plasma] (p = 1.00), and breastfeeding period [TM, 4.0 (4.0 – 11.5) months vs. NTM, 6.5 (4.0 – 18.8) months] (p = 0.53). This genetic protocol was approved by the Institutional Review Boards of the University of Florida, the Sabin Research Institute, and Children’s Hospital Los Angeles.

**Table 1.** Demographic, immune and viral characteristics of study subjects.

| Study group | Maternal |  |  |  |  | Infant age (days) |  |
| --- | --- | --- | --- | --- | --- | --- | --- |
| | Age (years) | Trimester | CD4 T cell (cells/μl) | Viral Load<sup>a</sup> | Duration of breastfeeding (months) | Negative PCR<sup>b</sup> | Positive PCR<sup>b</sup> |
| **TM1** | 22 | 2<sup>nd</sup> | 154 | 5.3 | 14 | 34 | 62 |
| **TM2** | 27 | 3<sup>rd</sup> | 110 | 5.5 | 4 | 35 | 63 |
| **TM3** | 33 | 3<sup>rd</sup> | 198 | 5.0 | 4 | 37 | 70 |
| **TM4** | 24 | 3<sup>rd</sup> | 137 | 4.9 | 4 | 28 | 63 |
| **NTM1** | 19 | 3<sup>rd</sup> | 181 | 5.3 | 4 | 738 |  |
| **NTM2** | 24 | 2<sup>nd</sup> | 115 | 5.0 | 22 | 731 |  |
| **NTM3** | 36 | 3<sup>rd</sup> | 223 | 5.3 | 4 | 730 |  |
| **NTM4** | 30 | 2<sup>nd</sup> | 246 | 5.1 | 9 | 639 |  |

a: Log<sub>10</sub> HIV-1 RNA copies/ml plasma; b: HIV-1 DNA

### Generation of amplicon library

Viral RNA was extracted from 280 μl of plasma using QIAamp Viral RNA Mini Kit (Qiagen, Valencia, CA). A library of HIV-1 *env* gp41 amplicons (318 nucleotides in length, including HR2, MPER, and 5’ MSD) was generated for each subject from 2,000 HIV-1 RNA copies by RT-PCR using SuperScript™ One-Step RT-PCR (Invitrogen, Carlsbad, California) followed by amplification using GoTaq colorless Master Mix (Promega, Madison, WI) (36). First round amplification used forward primer 251 (5’-GGG GCT GCT CTG GAA AAC TCA TCT-3’) and reverse primer 585 (5’-AAA CAC TAT ATG CTG AAA CAC CTA-3’) [nucleotides 8,011 - 8,035 and 8,345 - 8,469, respectively, in HIV-1_HXB2_ genome (37)], while second round amplification used forward A-257 (5’- <u>CGTATCGCCTCCCTCGCGCCATCAG</u> GCT CTG GAA AAC TCA TCT GCA CCA-3’) and reverse B-575 (5’-<u>CTATGCGCCTTGCCAGCCCGCTCAG</u> ATC CCT GCC TAA CTC TAT TCA CTA-3’) (positions 8,017 - 8,041 and 8,335 - 8,359, respectively) with adaptors A or B (underlined nucleotides in respective primer) incorporated at the 5’ ends. Amplicons were gel purified using QIAquick Gel Extraction Kit (Qiagen) as described (38), and submitted to the Interdisciplinary Center for Biotechnology Research at University of Florida for Titanium Amplicon 454-pyrosequencing reading from adaptor B using a Genome Sequencer FLX (454 Life Sciences) according to the manufacturer’s protocol.

### Sequence analysis

A bioinformatics pipeline was developed to facilitate analysis of large numbers of HIV-1 gp41 HR2-MPER-MSD sequence reads. The median (quartile range) number of raw reads was 56,647 (43,142 - 75,450) per subject. Sequences were submitted to NCBI public access database with accession numbers pending. A quality control step filtered a median (quartile range) of 7.5% (5.2% - 13.2%) low quality reads with ambiguous nucleotides, more than one error in either primer tag, or a length outside mean ± 2 SD length range, leaving median (quartile range) of 52,408 (37,541 - 71,533) quality sequences per sample. Depth of sequencing provided median (quartile range) of 27 (19 - 36)-fold coverage of input 2,000 HIV-1 RNA copies with no significant difference in sequence number or fold coverage among the samples between the groups. Quality MPER sequences were extracted from the entire HR2-MPER-MSD sequences by aligning to HIV-1_HXB2_ and to HIV-1 subtype C consensus sequence generated from HIV sequence database (39).

Nucleotide sequences were clustered at 3% genetic distance using ESPRIT (38,40,41) to develop a consensus sequence for each cluster that represents a sequence variant. Complexity of the HIV-1 population within each individual was evaluated by neighbor-joining (NJ) phylogenetic tree generated from consensus sequences with the maximum-likelihood composite model implemented in MEGA v5.2 (42,43). Statistical support was assessed by 1,000 bootstrap replicates. NJ trees were annotated manually in Adobe Illustrator CS4 (Adobe Systems Incorporated, San Jose, CA) to display frequencies of HIV-1 cluster variants. Frequencies of amino acid differences at each position compared to subtype B HIV-1_HXB2_ were calculated. Non-synonymous substitutions resulting in alteration of viral sensitivity to bn-HIV-Abs, including 2F5, 4E10, 10E8 and Z13e1, were identified by mapping to known resistant/sensitizing mutations (see Fig. S1) (21,29,44–60). Number and frequency of amino acid differences were compared between TM and NTM sequences. Positive selection at epitope-composing positions was inferred by Phylogenetic Analysis by Maximum Likelihood (PAML) (61). Hydropathy index and charge of each MPER consensus sequence were calculated using an in-house code (41,62,63).

Polymorphisms across all sequences were evaluated by biodiversity, expressed as operational taxonomic units (OTU), using rarefaction, while Chao1 algorithms in ESPRIT (40). Rarefaction curves display HIV-1 diversity over sequencing depth, and Chao1 infers maximum biodiversity within 2,000 input HIV-1 RNA copies (38,40,41).

### Statistical analysis

Groups were compared by unpaired t-test. Statistical analyses were performed using SAS version 9.1 (SAS Institute, Cary, NC) with P < 0.05 (two sided) defined as significant. Logistic regression was used to examine the effects of predicted hydropathy or charge of HIV-1 gp41 MPER and their interactions (exposures) on transmission (outcome).

## RESULTS

### Population structure

To evaluate the complexity of viral population structure within each individual, unrooted phylogenetic trees were constructed from consensus MPER sequence clusters. Overall, the analysis showed that sequences were correctly assigned to each individual with no sequence mixing among subjects. Within each subject HIV-1 populations were organized into one to three dominant clusters with thousands of sequences per cluster (Fig. 1). Dominant sequence clusters generally included a median (quartile range) of 47% (19% - 63%) of sequences. Sequences representing 0.25% to 10% of the viral population within an individual also appeared in low frequency (0 to 4) clusters surrounded by swarms of clusters with less abundant variants, usually representing < 0.25% of the population. The structure of viral populations based on gp41 regions was indistinguishable between TM and NTM and similar to HIV-1 populations based on gp120 V3 (38).

**Fig. 1.**
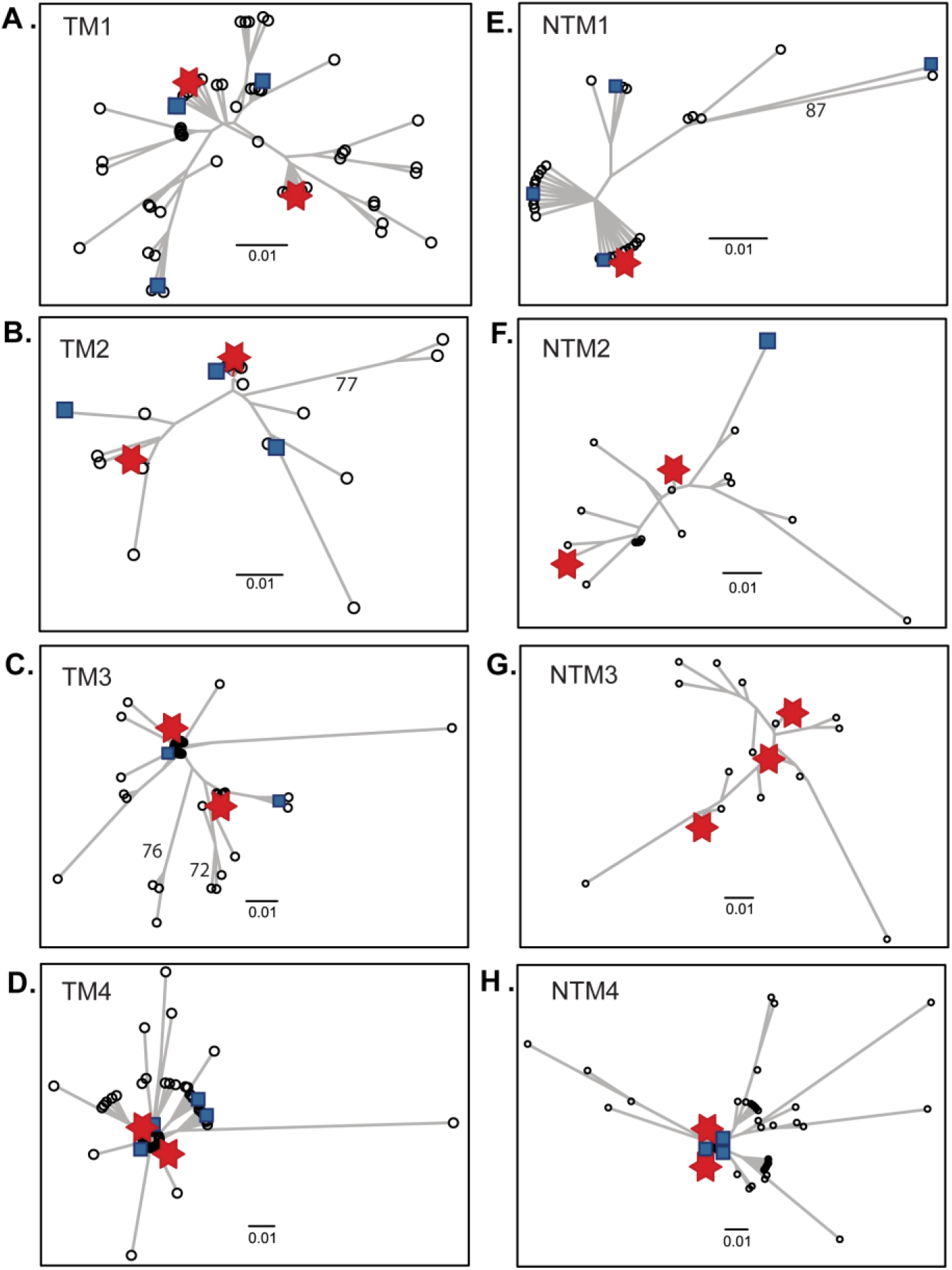
Organization of HIV-1 gp41 MPER populations. An unrooted neighbor-joining tree for each individual was developed from the deep sequencing data set clustered at 3% genetic distance. Each branch represents a consensus sequence of HIV-1 gp41 MPER within 3% genetic distance. Symbols represent the proportion of total deep sequences in a cluster: Ο, ≤ 0.25%; ■, > 0.25 % to 10%;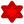, >10%.

### Biodiversity of HIV-1 MPER quasispecies

Biodiversity of HIV-1 MPER nucleotide sequences within each individual was assessed using rarefaction curves. HIV-1 MPER nucleotide sequences among TM displayed biodiversity ranging from 26 to 110 OTU, which was approximately 50% greater than biodiversity ranging from 18 to 77 OTU among NTM (Fig. 2A). When maximum biodiversity within 2,000 HIV-1 RNA copies was estimated, viral populations among TM, compared to populations among NTM, displayed a trend toward greater median maximum biodiversity [median (quartile range): 87 (66 - 160) OTU versus 33 (28 - 125) OTU, respectively, p = 0.33] (Fig. 2B).

**Fig. 2.**
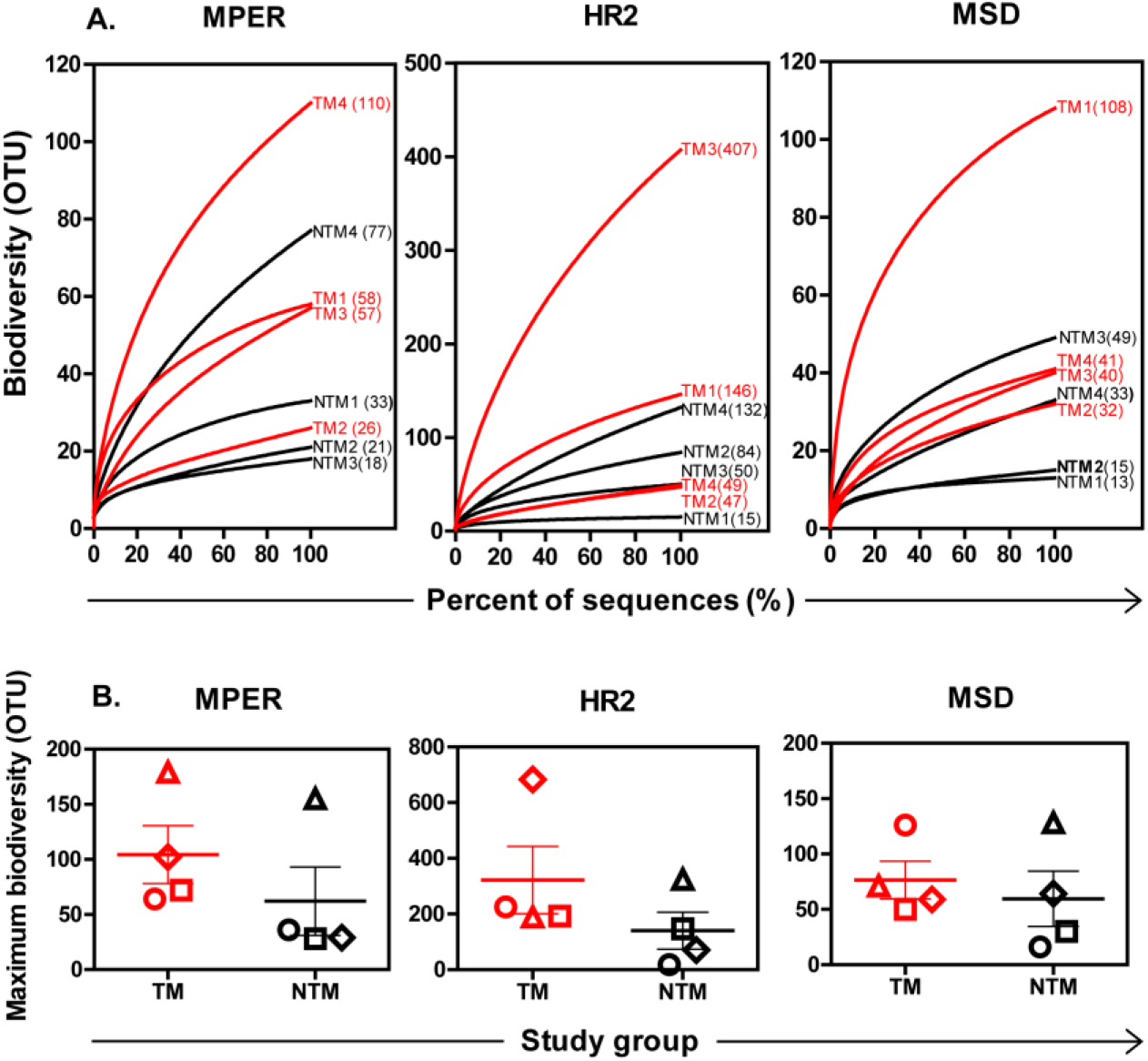
Biodiversity among HIV-1 viral populations. Nucleotide deep sequences of HIV-1 MPER (66 bp), or HR2 (108 bp), or MSD (63 bp) from each individual were clustered at 3% genetic distances and displayed as rarefaction curves **(A)** and Chao1 values **(B)**. **A.** Y-axis, number of OTU (number of sequence clusters); x-axis, percent of total deep sequences (sequences sampled ÷ total number of sequences x 100%). Rarefaction curves show HIV-1 variants from TMs (red) or NTMs (black), respectively. Numbers of OTU at the end of curves represent biodiversity calculated from rarefaction curve at the sequence depth (100% of deep sequences). **B.**Y-axis, maximum number of OTU within 2,000 input viral copies estimated by Chao1 algorithm based on rarefaction curve of HIV-1 variants from each subjects (40); x-axis, study group, TM or NTM, respectively. Symbols: Ο, subject #1; □, subject #2; ◊, subject #3; Δ, subject #4. Red symbols, TM; black symbols, NTM

To determine if differences in biodiversity between TM and NTM were restricted to MPER or extended to adjacent regions in gp41, similar analyses were applied to HR2 and to MSD sequences (Fig. 2). Overall, mean estimated maximum biodiversity was more than 2-fold greater in HR2 than in MPER among TM or NTM groups, reflecting in part that the HR2 region (102 nucleotides) is almost twice as long as MPER (66 nucleotides). MSD encoding regions are similar to MPER in length and displayed similar biodiversity between NTM and TM groups, although maximum biodiversity in MSD compared to MPER was reduced among TM group (Fig. 2B).

### Amino acid substitutions in HIV-1 MPER

Biodiversity evaluated at the nucleotide sequence level was reflected in diversity among amino acid residues in MPER (Fig. 3), as well as in HR2 and in MSD regions (see Figs. S2 and S3), indicating that a preponderance of nucleotide polymorphisms within each region involved nonsynonymous changes. HIV-1 MPER variants among TM had changes at more amino acid positions than NTM [median (quartile range) 14 (12 - 16) vs. 9 (8 - 14) positions per person, respectively], with amino acid changes in six positions (663, 666, 672, 673, 680 and 681) observed exclusively in TM viral populations. HIV-1 MPER variants from TM also had more amino acid substitutions per position than NTM [median (quartile range): 7 (4 - 9) vs. 3 (1 - 7) respectively, p = 0.04]. While the MPER reference sequence for subtype B includes a single N-linked glycosylation motif (positions 674 to 676), the subtype C consensus MPER sequence lacks a similar motif. Although some polymorphisms at position 674 would introduce a motif at low frequency, the number of N-linked glycosylation motifs in MPER was similar among viral populations from TM and NTM. MPER amino acid residues under positive selection were limited (N674G and K683R in TM1, S668K in TM4, N677R in NTM2, and K665R, T676S and K683R in NTM4) with no significant difference between TM and NTM (Fig. 3).

**Fig. 3.**
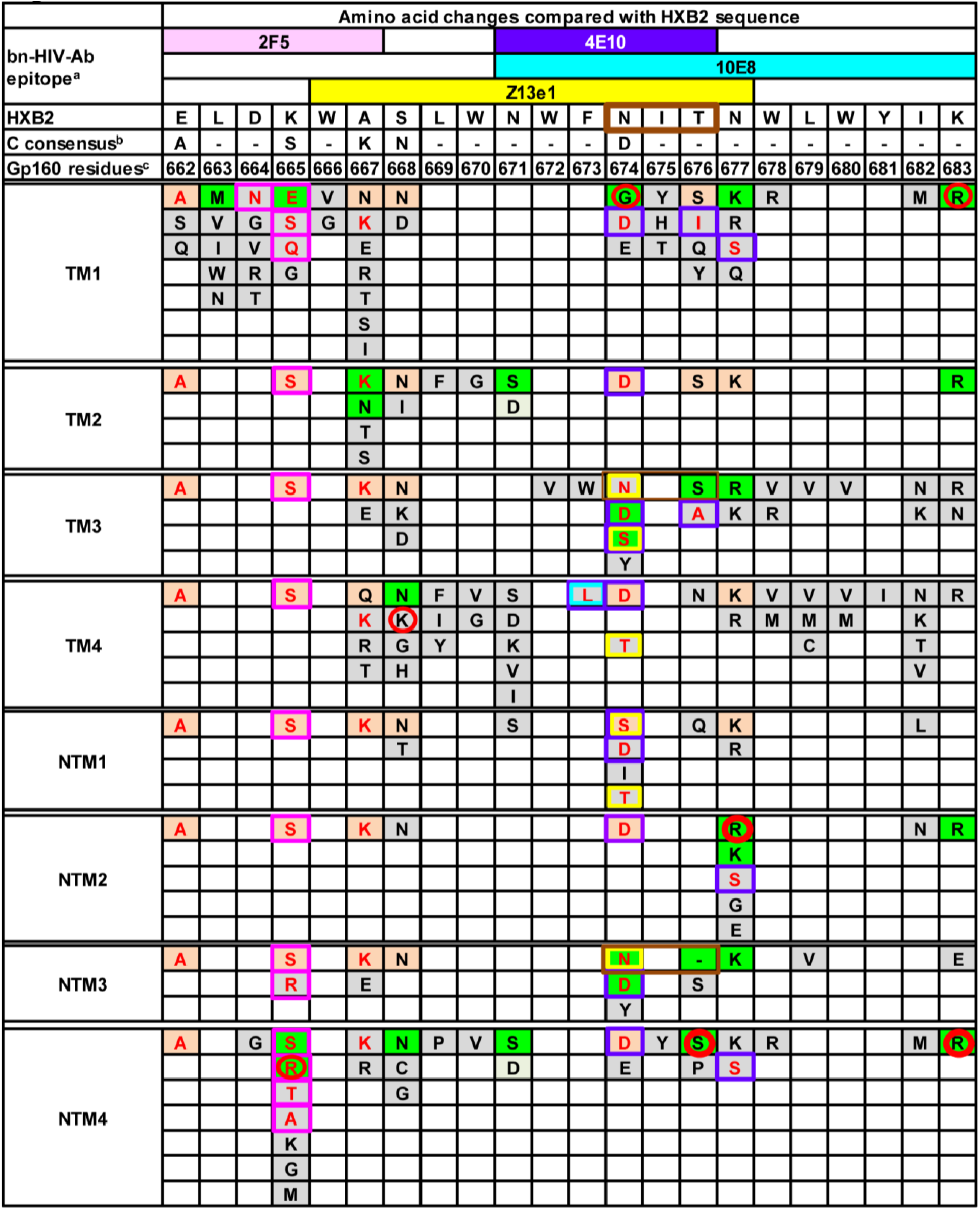
Amino acid changes in MPER compared with HXB2 sequence. Amino acid residue (a single letter code) which differs from HXB2 sequence was shown in each space with **red letter** representing amino acid residue resistant to bn-HIV-Ab(s) and black letter depicting amino acid with unknown effect on bn-HIV-Ab susceptibility. Color scheme is used to define frequency of amino acid substitution with **beige** representing residues in > 80% of HIV-1 MPER variants; **green** depicting residues in > 10% to 80% of HIV-1 MPER variants; and **grey** representing residues in < 1% to 10% of HIV-1 MPER variants. Substitutions outlined in **pink** are resistant to 2F5; **purple** are resistant to 4E10; **blue** are resistant to 10E8; and **yellow** are resistant to Z13e1. Residues under positive selection are circled by **red**. N-linked glycosylation motifs (NXS/T) are outlined by **dark brown**. a: epitope reported in HIV molecular immunology database (91); b: subtype C consensus sequence generated from HIV sequence database (39); c: gp160 amino acid residues 662 to 683 are residues 151 to 172 in gp41 (37); dash (-): amino acid identity between HIV-1_HXB2_ and subtype C consensus.

### Changes in antibody response epitopes in MPER

Amino acid substitutions in MPER epitopes might alter susceptibility (i.e., sensitivity or resistance) to bn-HIV-Abs, including 2F5, 4E10, 10E8 and Z13e1 (see Fig. S1). A bioinformatics approach was applied to evaluate a potential impact on neutralization susceptibility by amino acid polymorphisms in MPER among the sequences. Overall, the neutralization effects by many of the MPER polymorphisms identified by deep sequencing were undefined (Fig. 3), although no known sensitizing variants, even at low frequency, were identified in any subject (44,46,50,53,64). In contrast, some MPER polymorphisms were predicted to be associated with resistance to neutralization by 2F5, Z13e1, 4E10, or 10E8 (21,29,45,47–49,51–55,57,58,60,65–67). For example, all subjects harbored dominant virus populations with known subtype C amino substitutions E662A, K665S and A667K conferring 2F5 resistance (48,53,65–67). Additional 2F5 resistant polymorphisms D664N and K665E/Q/R/T/A (45,47,48,51,54,60,68) were identified in 3 individuals (TM1, NTM3, and NTM4). At least one of four 4E10 resistant substitutions (F673L, N674D/S, T676/I/A, or N677S) was identified in each individual (29,49,50,52,54,55,58). Resistance substitutions to Z13e1 (D674N/S/T) (57) appeared in several TM (TM3 and TM4) and NTM (NTM1 and NTM3), while 10E8 resistant mutation F673L (52) was observed only in TM4. Overall, polymorphic substitutions with predicted resistance phenotypes were identified with variable frequency in most individuals independent of transmission outcomes.

### Distinct biochemical characteristics of HIV-1 MPER populations between TM and NTM

To evaluate whether or not predicted amino acid substitutions might alter the biochemical features of MPER, distribution of hydropathy or charge at the population level within TM or NTM MPER was assessed (Fig. 4A). TM viral populations compared with NTM demonstrated a left-shift towards increased frequencies of hydrophilic MPER variants with a median (quartile range, QR) hydropathy index of −10 (QR, −12.5 to −9.6), significantly lower than NTM variants with a median of −7.3 (QR, −10.4 to −5.1) (p <0.0001). The difference in hydropathy index between TM and NTM was concentrated among variants that appeared with reduced frequency (≤ 20%) (P < 0.0001), but not among high frequency variants (> 20%) (p = 0.34). Low-frequency variants were uniquely identified by NGS, and not found when clonal or single genome sequences were analyzed (29) (data not shown). When charge of MPER amino acids was assessed, a clear right-shift towards an increase in frequencies of MPER variants with greater positive charges occurred in TM with significantly greater net charges (median 2.0; QR, 1.0 to 2.0) compared with NTM (median 1.0; QR, 1.0 to 2.0) (p<0.001) (Fig. 4B). The distinct differences in biochemical features between TM and NTM gp41 populations were restricted to MPER and failed to extend into flanking HR2 or MSD domains (Fig. 4).

**Fig. 4.**
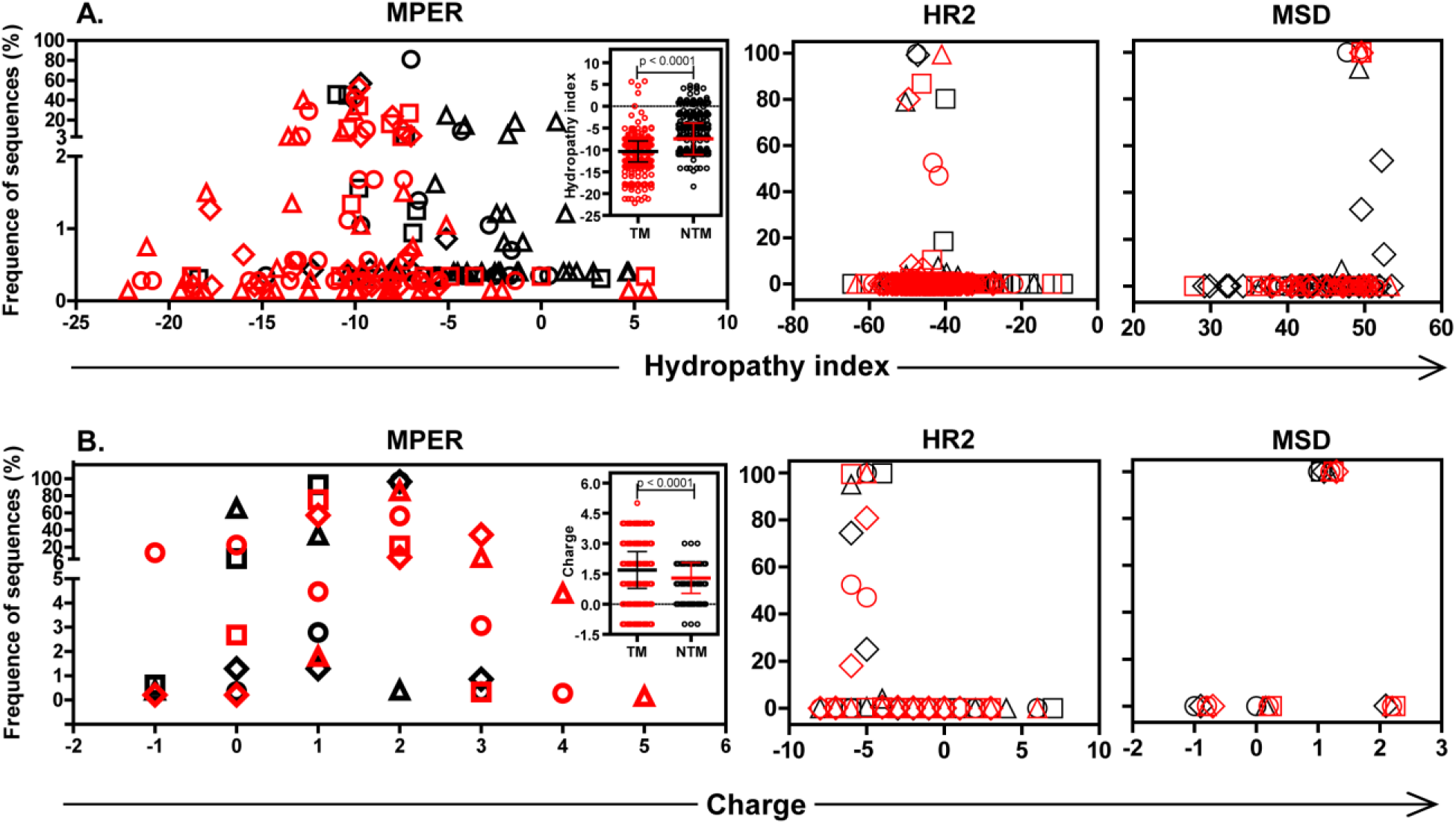
Biochemical characteristics of HIV-1 viral variants. Frequency distribution of **A.** hydropathy indexes with each symbol representing the percent of consensus sequences with that particular hydropathy index, or **B.** net charge of HIV-1 viral variants with each symbol depicting percent of consensus sequences with that particular net charge of MPER, HR2 or MSD from TMs (red symbols) or NTMs (black symbols). Symbols: Ο, subject #1; □, subject #2; ◊, subject #3; Δ, subject #4. Inserts in **A**and **B**show significantly lower hydropathy index and significantly higher net charge respectively in HIV-1 MPER variants from TM in contrast to NTM with each point representing hydropathy index **(A)** or net charge **(B)** of each consensus MPER sequence.

Logistic regression analysis indicated that an increase in MPER hydrophobicity was significantly associated with reduced odds of transmission by breast feeding (p < 0.0001), while positive charged MPER regions showed a close relationship with breast milk transmission (p < 0.0001). Logistic regression statistics revealed a significant interactive effect on transmission between hydropathy and charge (p<0.0001). For negative, neutral or positive charged regions, odds ratios were 0.741, 0.416 and 0.781 respectively for a one-unit increase in net hydropathy (95% confidence interval 0.738 - 0.744, 0.413 - 0.419 and 0.699 - 0.873, respectively). Charge has an opposite effect on transmission for negative and positive hydropathy. Increase of net charge was significantly associated with reduced odds of transmission for negative hydropathy (OR = 0.627, 95% CI, 0.622 - 0.632), while for positive hydropathy, net charge increase was significantly associated with elevated odds of transmission (OR = 6.358, 95% CI, 3.772 - 10.718).

## DISCUSSION

Breast milk is essential for infant development and health particularly in resource limited settings (69–72). Unfortunately breast feeding remains a major source of global pediatric HIV-1 infection reflecting, in part, limited parameters to identify women at high risk for viral transmission by breastfeeding and the challenges of providing therapeutic interventions for the duration of the breast feeding period (73–76). HIV-1 variants that establish new infections by breastfeeding generally occur at low frequency in the transmitting viral population, are characterized by shorter and underglycosylated gp120 Envelopes, and may represent escape from neutralizing antibodies targeting epitopes in both gp120 and gp41 MPER (9–12,77). Our exploratory studies of HIV-1 variants by metagenomic approaches identified distinct features of gestational MPER populations that distinguished between women who did or did not subsequently transmit during breastfeeding. Transmission outcome groups in our study were well balanced in age, plasma viral load, CD4 T-cell counts and breastfeeding practices, which in combination with the depth of sequencing from each individual provided statistical sensitivity. As anticipated virus populations in plasma during pregnancy among women who subsequently transmitted via breastfeeding displayed greater biodiversity. A higher frequency of HIV-1 MPER variants with hydrophilic and positively charged amino acid residues among TM compared with NTM was discovered. The characteristics could only be evaluated at the population level by NGS, as conventional clonal sequencing biases the population towards dominant variants. Phenotypic differences in peripheral blood viral populations overtime that related to subsequent transmission were evident by the third trimester of pregnancy about the time of lactogenesis (27). While our current study was designed as a cross sectional comparison of maternal virus populations during gestation, whether or not biochemical differences among maternal viral populations present during pregnancy persist during breastfeeding and are related to infecting cell-free or cell-associated viruses in nursing babies are important questions for subsequent studies (78).

Positive selection for any single amino acid change was limited, as was modulation of glycan motifs across MPER. Sensitivity to bn-HIV-Ab, either alone or in combinations, by the novel amino acids in each MPER allele within an individual is difficult to predict with complete accuracy, may differ by subtype (77) and necessitates direct assessment for neutralization susceptibility (79). Absence of clear bn-HIV-Ab resistance genotypic profiles during pregnancy that distinguish between TM and NTM does not rule out a subsequent role for neutralization resistance in MTCT by breast milk. Yet, polymorphic amino acid positions within MPER during pregnancy frequently mapped outside motifs associated with known bn-HIV-Ab, raising the possibility that factors other than antibody selection contribute to the differences in MPER characteristics between TM and NTM. For example, a significant role in membrane fusion played by MPER requires functional assays to evaluate the consequences by biochemical variants of MPER for viral entry into different host cells or for crossing mucosal barriers.

HIV-1 gp41 MPER plays a critical role in HIV-1 fusion by perturbing the architecture of the bilayer envelope (80–83). Distribution of hydrophobic amino acid in MPER can modulate membrane fusion (80,84). Electrostatic interaction between viral particle and negatively charged lipid membrane may also play a role in viral entry (85). Logistic regression analysis indicated an interactive effect of hydropathy and charge of HIV-1 MPER variants on breast milk transmission outcome in our study. Similar to our study of gp41 MPER, a significant difference in hydropathy in gp120 between TM and NTM in intrauterine transmission was reported in another study (86), suggesting that intrauterine transmission is associated with maternal envelope quasispecies with altered cellular tropism or affinity for coreceptor molecules expressed on cells localized in the placenta. Together, both studies raise the possibility that antibody-independent mechanisms might contribute to transmission.

A novel aspect of our study is that differences in MPER were compared to flanking regions in gp41. While MPER regions displayed a trend toward increased maximum biodiversity, the striking biochemical characteristics of viral populations associated with MTCT by breastfeeding were restricted to MPER. Although HR2 and MSD segments that flank MPER were diverse, patterns of diversity were unrelated to transmission outcomes, perhaps reflecting HR2 interactions with HR1 or a role for MSD in anchoring gp41 in membranes (64,87–90). Overall, deep sequencing coupled with an efficient bioinformatics pipeline provided unprecedented coverage of HIV-1 gp41 MPER quasispecies combined with sensitive detection of low frequency variants that can only be captured by high coverage of input viral copies. Low frequency variants within viral populations are particularly critical and clinically relevant as transmitting viruses. Our proof of principle studies indicating that detailed characteristics of viral quasispecies months before transmission relate to transmission outcomes raises the possibility for predicting MTCT by breastfeeding and identifying mothers with high-risk viral populations.

## ACKNOWLEDGEMENTS

The authors wish to thank the study volunteers for participating. Research was supported in part by Elizabeth Glaser Pediatric AIDS Foundation (GMA and MMG); NIH/NIAID R01 AI065265 (MMG); Pediatric Immune Deficiency Center, University of Florida, College of Medicine (MMG); and Stephany W. Holloway University Chair for AIDS Research (MMG); NIH funds through an International Maternal Pediatric Adolescent AIDS Clinical Trials Group (IMPAACT) Virology Developmental Laboratory award (GMA) (UM1AI106716). Overall support for IMPAACT was provided by the National Institute of Allergy and Infectious Diseases (NIAID) of the National Institutes of Health (NIH) under Award Numbers UM1AI068632 (IMPAACT LOC), UM1AI068616 (IMPAACT SDMC) and UM1AI106716 (IMPAACT LC), with co-funding from the Eunice Kennedy Shriver National Institute of Child Health and Human Development (NICHD) and the National Institute of Mental Health (NIMH). The content is solely the responsibility of the authors and does not necessarily represent the official views of the NIH.

## SUPPLEMENTAL FIGURES

**Fig. S1.**
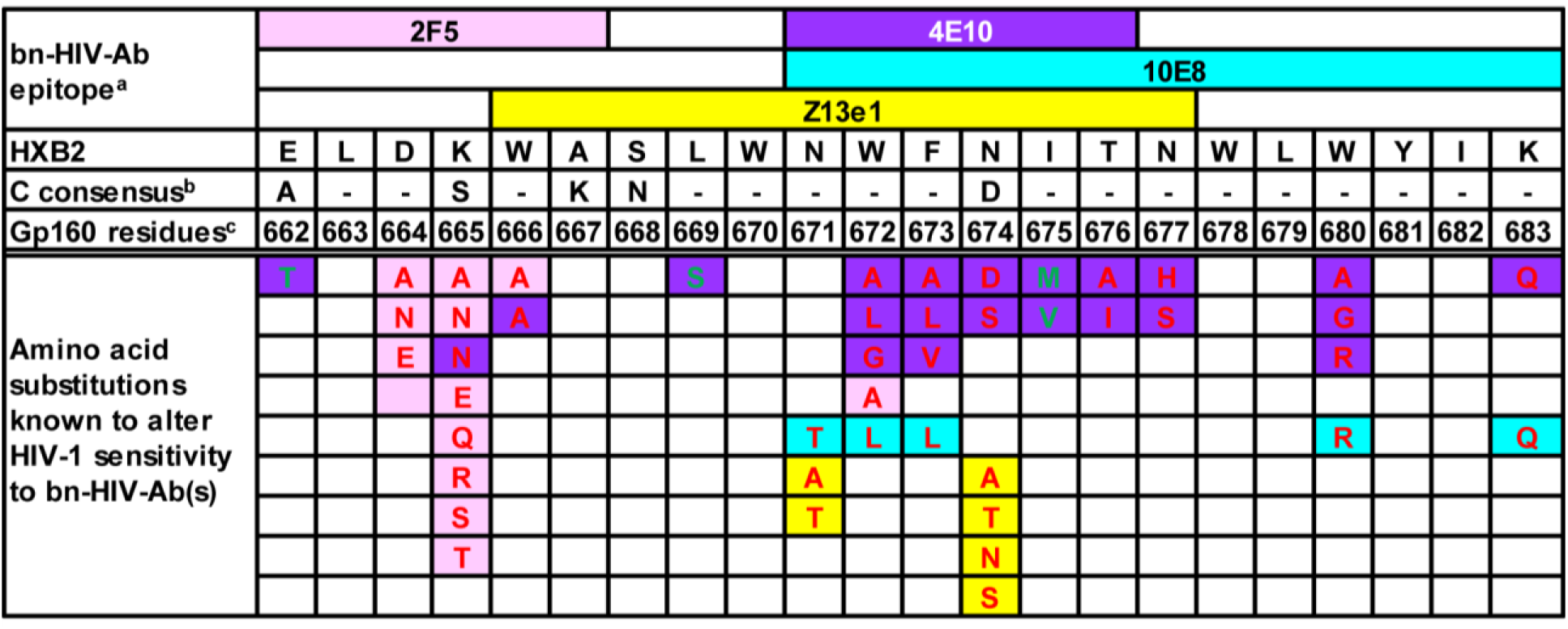
Amino acid changes known to alter HIV-1 sensitivity to bn-HIV-Ab(s). Amino acid residue (a single letter code) known to alter HIV-1 sensitivity to bn-HIV-Ab(s) is shown in each space with **red letter** representing Ab-resistant amino acid substitution (1–15), and green letter depicting amino acid substitution increasing sensitivity to Ab-neutralization (5,16–19). Colors code epitopes of known HIV-1 MPER antibodies, and correspondent resistant/sensitizing amino acid residues with **pink** for 2F5, **purple** for 4E10, **blue** for 10E8 and **yellow** for Z13e1. a: epitope reported in HIV molecular immunology database (20); b: subtype C consensus sequence generated from HIV sequence database {Los Alamos National Laboratory, 14 A.D. 73 /id}; c: gp160 amino acid residues 662 to 683 are residues 151 to 172 in gp41 (21); dash (-): amino acid identity between HIV-1_HXB2_ and subtype C consensus.

**Fig. S2.**
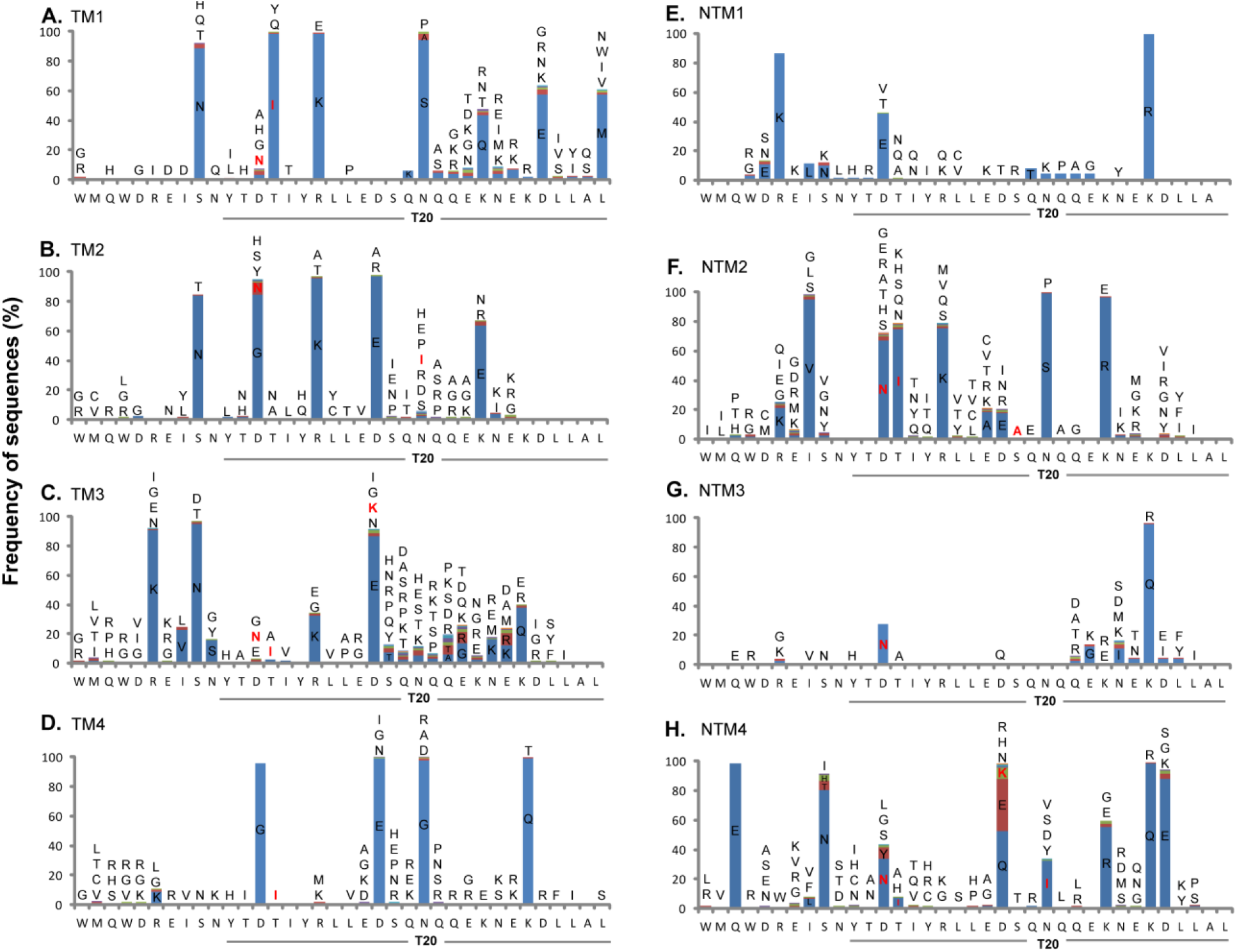
Amino acid changes in HR2 compared with HXB2 sequence. Amino acid residue(s) (single letter code) differing from HXB2 sequence is/are illustrated at each position of HR2 with **red letter** representing T20-resistant mutation (22–24).

**Fig. S3.**
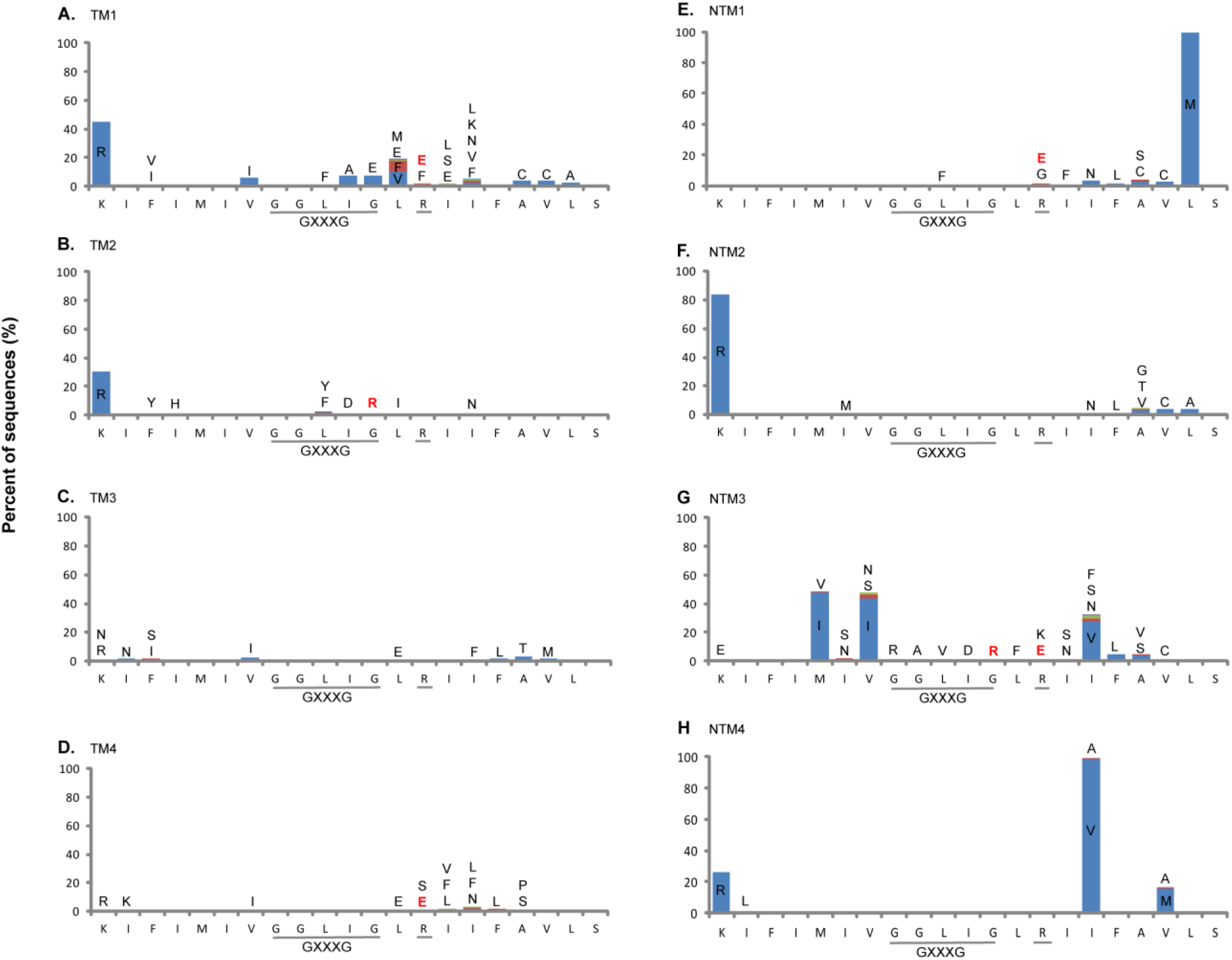
Amino acid changes in MSD compared with HXB2 sequence. Amino acid residue(s) (single letter code) differing from HXB2 sequence is/are illustrated at each position of MSD with GXXXG motif and R696 underlined due to their important roles in membrane fusion and fusion abolishing mutation G694/R or R696/E/D highlighted in red (25–27).

## REFERENCES

1. Dabis F, Msellati P, Newell ML, Halsey N, Van de Perre P, Peckham C, Lepage P. 1995. Methodology of intervention trials to reduce mother to child transmission of HIV with special reference to developing countries. International Working Group on Mother to Child Transmission of HIV. AIDS 9 Suppl A:S67–S74.

2. Pan E, Wara D, DeCarlo P, Freedman B. Is Mother-to-Child HIV Transmission Preventable? 2002. http://caps.ucsf.edu/archives/factsheets/mother-to-child-transmission-mtct

3. Volmink J, Siegfried NL, van der Merwe L, Brocklehurst P. 2007. Antiretrovirals for reducing the risk of mother-to-child transmission of HIV infection. Cochrane Database Syst Rev 24:CD003510. doi:10.1002/14651858.CD003510.pub2 [doi].

4. World Health Organization. Mother-to-child transmission of HIV. 2014. http://www.who.int/hiv/topics/mtct/en/

5. Govender T, Coovadia H. 2014. Eliminating mother to child transmission of HIV-1 and keeping mothers alive: recent progress. J Infect 68 Suppl 1:S57–S62. doi:S0163-4453(13)00282-X [pii];10.1016/j.jinf.2013.09.015 [doi].

6. Taha TE, Kumwenda J, Cole SR, Hoover DR, Kafulafula G, Fowler MG, Thigpen MC, Li Q, Kumwenda NI, Mofenson L. 2009. Postnatal HIV-1 transmission after cessation of infant extended antiretroviral prophylaxis and effect of maternal highly active antiretroviral therapy. J Infect Dis 200:1490–1497. doi:10.1086/644598 [doi].

7. Mofenson LM. 2010. Antiretroviral drugs to prevent breastfeeding HIV transmission. Antivir Ther 15:537–553. doi:10.3851/IMP1574 [doi].

8. Rainwater SM, Wu X, Nduati R, Nedellec R, Mosier D, John-Stewart G, Mbori-Ngacha D, Overbaugh J. 2007. Cloning and characterization of functional subtype A HIV-1 envelope variants transmitted through breastfeeding. Curr HIV Res 5:189–197. doi:10.2174/157016207780076986#sthash.2eLyecCO.dpuf.

9. Russell ES, Kwiek JJ, Keys J, Barton K, Mwapasa V, Montefiori DC, Meshnick SR, Swanstrom R. 2011. The genetic bottleneck in vertical transmission of subtype C HIV-1 is not driven by selection of especially neutralization-resistant virus from the maternal viral population. J Virol 85:8253–8262. doi:JVI.00197-11 [pii];10.1128/JVI.00197-11 [doi].

10. Russell ES, Ojeda S, Fouda GG, Meshnick SR, Montefiori D, Permar SR, Swanstrom R. 2013. Short communication: HIV type 1 subtype C variants transmitted through the bottleneck of breastfeeding are sensitive to new generation broadly neutralizing antibodies directed against quaternary and CD4-binding site epitopes. AIDS Res Hum Retroviruses 29:511–515. doi:10.1089/AID.2012.0197 [doi].

11. Wolinsky SM, Wike CM, Korber BT, Hutto C, Parks WP, Rosenblum LL, Kunstman KJ, Furtado MR, Munoz JL. 1992. Selective transmission of human immunodeficiency virus type-1 variants from mothers to infants. Science 255:1134–1137.

12. Zhang H, Tully DC, Hoffmann FG, He J, Kankasa C, Wood C. 2010. Restricted genetic diversity of HIV-1 subtype C envelope glycoprotein from perinatally infected Zambian infants. PLoS One 5:e9294. doi:10.1371/journal.pone.0009294 [doi].

13. Boeras DI, Hraber PT, Hurlston M, Evans-Strickfaden T, Bhattacharya T, Giorgi EE, Mulenga J, Karita E, Korber BT, Allen S, Hart CE, Derdeyn CA, Hunter E. 2011. Role of donor genital tract HIV-1 diversity in the transmission bottleneck. Proc Natl Acad Sci U S A 108:E1156–E1163. doi:1103764108 [pii];10.1073/pnas.1103764108 [doi].

14. Chohan B, Lang D, Sagar M, Korber B, Lavreys L, Richardson B, Overbaugh J. 2005. Selection for human immunodeficiency virus type 1 envelope glycosylation variants with shorter V1-V2 loop sequences occurs during transmission of certain genetic subtypes and may impact viral RNA levels. J Virol 79:6528–6531. doi:79/10/6528 [pii];10.1128/JVI.79.10.6528-6531.2005 [doi].

15. Derdeyn CA, Decker JM, Bibollet-Ruche F, Mokili JL, Muldoon M, Denham SA, Heil ML, Kasolo F, Musonda R, Hahn BH, Shaw GM, Korber BT, Allen S, Hunter E. 2004. Envelope-constrained neutralization-sensitive HIV-1 after heterosexual transmission. Science 303:2019–2022. doi:10.1126/science.1093137 [doi];303/5666/2019 [pii].

16. Salazar-Gonzalez JF, Bailes E, Pham KT, Salazar MG, Guffey MB, Keele BF, Derdeyn CA, Farmer P, Hunter E, Allen S, Manigart O, Mulenga J, Anderson JA, Swanstrom R, Haynes BF, Athreya GS, Korber BT, Sharp PM, Shaw GM, Hahn BH. 2008. Deciphering human immunodeficiency virus type 1 transmission and early envelope diversification by single-genome amplification and sequencing. J Virol 82:3952–3970. doi:JVI.02660-07 [pii];10.1128/JVI.02660-07 [doi].

17. Burton DR, Stanfield RL, Wilson IA. 2005. Antibody vs. HIV in a clash of evolutionary titans. Proc Natl Acad Sci U S A 102:14943–14948. doi:0505126102 [pii];10.1073/pnas.0505126102 [doi].

18. Douek DC, Kwong PD, Nabel GJ. 2006. The rational design of an AIDS vaccine. Cell 124:677–681. doi:S0092-8674(06)00180-2 [pii];10.1016/j.cell.2006.02.005 [doi].

19. Montefiori DC. 2005. Neutralizing antibodies take a swipe at HIV in vivo. Nat Med 11:593–594. doi:nm0605-593 [pii];10.1038/nm0605-593 [doi].

20. Montero M, van Houten NE, Wang X, Scott JK. 2008. The membrane-proximal external region of the human immunodeficiency virus type 1 envelope: dominant site of antibody neutralization and target for vaccine design. Microbiol Mol Biol Rev 72:54–84, table. doi:72/1/54 [pii];10.1128/MMBR.00020-07 [doi].

21. Zwick MB, Jensen R, Church S, Wang M, Stiegler G, Kunert R, Katinger H, Burton DR. 2005. Anti-human immunodeficiency virus type 1 (HIV-1) antibodies 2F5 and 4E10 require surprisingly few crucial residues in the membrane-proximal external region of glycoprotein gp41 to neutralize HIV-1. J Virol 79:1252–1261. doi:79/2/1252 [pii];10.1128/JVI.79.2.1252-1261.2005 [doi].

22. Baan E, de RA, Stax M, Sanders RW, Luchters S, Vyankandondera J, Lange JM, Pollakis G, Paxton WA. 2013. HIV-1 autologous antibody neutralization associates with mother to child transmission. PLoS One 8:e69274. doi:10.1371/journal.pone.0069274 [doi];PONE-D-13-14798 [pii].

23. Diomede L, Nyoka S, Pastori C, Scotti L, Zambon A, Sherman G, Gray CM, Sarzotti-Kelsoe M, Lopalco L. 2012. Passively transmitted gp41 antibodies in babies born from HIV-1 subtype C-seropositive women: correlation between fine specificity and protection. J Virol 86:4129–4138. doi:JVI.06359-11 [pii];10.1128/JVI.06359-11 [doi].

24. Guevara H, Casseb J, Zijenah LS, Mbizvo M, Oceguera LF, III, Hanson CV, Katzenstein DA, Hendry RM. 2002. Maternal HIV-1 antibody and vertical transmission in subtype C virus infection. J Acquir Immune Defic Syndr 29:435–440.

25. Lallemant M, Baillou A, Lallemant-Le CS, Nzingoula S, Mampaka M, M'Pele P, Barin F, Essex M. 1994. Maternal antibody response at delivery and perinatal transmission of human immunodeficiency virus type 1 in African women. Lancet 343:1001–1005. doi:S0140-6736(94)90126-0 [pii].

26. Pancino G, Leste-Lasserre T, Burgard M, Costagliola D, Ivanoff S, Blanche S, Rouzioux C, Sonigo P. 1998. Apparent enhancement of perinatal transmission of human immunodeficiency virus type 1 by high maternal anti-gp160 antibody titer. J Infect Dis 177:1737–1741.

27. Gray RR, Salemi M, Lowe A, Nakamura KJ, Decker WD, Sinkala M, Kankasa C, Mulligan CJ, Thea DM, Kuhn L, Aldrovandi G, Goodenow MM. 2011. Multiple independent lineages of HIV-1 persist in breast milk and plasma. AIDS 25:143–152. doi:10.1097/QAD.0b013e328340fdaf [doi];00002030-201101140-00003 [pii].

28. Kuhn L, Aldrovandi GM, Sinkala M, Kankasa C, Semrau K, Mwiya M, Kasonde P, Scott N, Vwalika C, Walter J, Bulterys M, Tsai WY, Thea DM. 2008. Effects of early, abrupt weaning on HIV-free survival of children in Zambia. N Engl J Med 359:130–141. doi:NEJMoa073788 [pii];10.1056/NEJMoa073788 [doi].

29. Nakamura KJ, Gach JS, Jones L, Semrau K, Walter J, Bibollet-Ruche F, Decker JM, Heath L, Decker WD, Sinkala M, Kankasa C, Thea D, Mullins J, Kuhn L, Zwick MB, Aldrovandi GM. 2010. 4E10-resistant HIV-1 isolated from four subjects with rare membrane-proximal external region polymorphisms. PLoS One 5:e9786. doi:10.1371/journal.pone.0009786 [doi].

30. Becquart P, Chomont N, Roques P, Ayouba A, Kazatchkine MD, Belec L, Hocini H. 2002. Compartmentalization of HIV-1 between breast milk and blood of HIV-infected mothers. Virology 300:109–117. doi:S0042682202915370 [pii].

31. Gantt S, Carlsson J, Heath L, Bull ME, Shetty AK, Mutsvangwa J, Musingwini G, Woelk G, Zijenah LS, Katzenstein DA, Mullins JI, Frenkel LM. 2010. Genetic analyses of HIV-1 env sequences demonstrate limited compartmentalization in breast milk and suggest viral replication within the breast that increases with mastitis. J Virol 84:10812–10819. doi:JVI.00543-10 [pii];10.1128/JVI.00543-10 [doi].

32. Heath L, Conway S, Jones L, Semrau K, Nakamura K, Walter J, Decker WD, Hong J, Chen T, Heil M, Sinkala M, Kankasa C, Thea DM, Kuhn L, Mullins JI, Aldrovandi GM. 2010. Restriction of HIV-1 genotypes in breast milk does not account for the population transmission genetic bottleneck that occurs following transmission. PLoS One 5:e10213. doi:10.1371/journal.pone.0010213 [doi].

33. Henderson GJ, Hoffman NG, Ping LH, Fiscus SA, Hoffman IF, Kitrinos KM, Banda T, Martinson FE, Kazembe PN, Chilongozi DA, Cohen MS, Swanstrom R. 2004. HIV-1 populations in blood and breast milk are similar. Virology 330:295–303. doi:S0042-6822(04)00591-4 [pii];10.1016/j.virol.2004.09.004 [doi].

34. Salazar-Gonzalez JF, Salazar MG, Learn GH, Fouda GG, Kang HH, Mahlokozera T, Wilks AB, Lovingood RV, Stacey A, Kalilani L, Meshnick SR, Borrow P, Montefiori DC, Denny TN, Letvin NL, Shaw GM, Hahn BH, Permar SR. 2011. Origin and evolution of HIV-1 in breast milk determined by single-genome amplification and sequencing. J Virol 85:2751–2763. doi:JVI.02316-10 [pii];10.1128/JVI.02316-10 [doi].

35. Becquart P, Courgnaud V, Willumsen J, Van de Perre P. 2007. Diversity of HIV-1 RNA and DNA in breast milk from HIV-1-infected mothers. Virology 363:256–260. doi:S0042-6822(07)00088-8 [pii];10.1016/j.virol.2007.02.003 [doi].

36. Coberley CR, Kohler JJ, Brown JN, Oshier JT, Baker HV, Popp MP, Sleasman JW, Goodenow MM. 2004. Impact on genetic networks in human macrophages by a CCR5 strain of human immunodeficiency virus type 1. J Virol 78:11477–11486. doi:78/21/11477 [pii];10.1128/JVI.78.21.11477-11486.2004 [doi].

37. Ratner L, Haseltine W, Patarca R, Livak KJ, Starcich B, Josephs SF, Doran ER, Rafalski JA, Whitehorn EA, Baumeister K,. 1985. Complete nucleotide sequence of the AIDS virus, HTLV-III. Nature 313:277–284.

38. Yin L, Liu L, Sun Y, Hou W, Lowe AC, Gardner BP, Salemi M, Williams WB, Farmerie WG, Sleasman JW, Goodenow MM. 2012. High-resolution deep sequencing reveals biodiversity, population structure, and persistence of HIV-1 quasispecies within host ecosystems. Retrovirology 9:108. doi:1742-4690-9-108 [pii];10.1186/1742-4690-9-108 [doi].

39. Los Alamos National Laboratory. HIV sequence database. 10-23-0014. http://www.hiv.lanl.gov/content/sequence/HIV/mainpage.html

40. Sun Y, Cai Y, Liu L, Yu F, Farrell ML, McKendree W, Farmerie W. 2009. ESPRIT: estimating species richness using large collections of 16S rRNA pyrosequences. Nucleic Acids Res 37:e76. doi:gkp285 [pii];10.1093/nar/gkp285 [doi].

41. Yin L, Hou W, Liu L, Cai Y, Wallet MA, Gardner BP, Chang K, Lowe AC, Rodriguez CA, Sriaroon P, Farmerie WG, Sleasman JW, Goodenow MM. 2013. IgM Repertoire Biodiversity is Reduced in HIV-1 Infection and Systemic Lupus Erythematosus. Front Immunol 4:373. doi:10.3389/fimmu.2013.00373 [doi].

42. Koichiro Tamura, Glen Stecher Daniel Peterson Sudhir Kumar. MEGA molecular revolutionary genetics analysis. 2014. http://www.megasoftware.net/

43. Tamura K, Peterson D, Peterson N, Stecher G, Nei M, Kumar S. 2011. MEGA5: molecular evolutionary genetics analysis using maximum likelihood, evolutionary distance, and maximum parsimony methods. Mol Biol Evol 28:2731–2739. doi:msr121 [pii];10.1093/molbev/msr121 [doi].

44. Blish CA, Nguyen MA, Overbaugh J. 2008. Enhancing exposure of HIV-1 neutralization epitopes through mutations in gp41. PLoS Med 5:e9. doi:07-PLME-RA-0379 [pii];10.1371/journal.pmed.0050009 [doi].

45. Bunnik EM, van Gils MJ, Lobbrecht MS, Pisas L, van Nuenen AC, Schuitemaker H. 2009. Changing sensitivity to broadly neutralizing antibodies b12, 2G12, 2F5, and 4E10 of primary subtype B human immunodeficiency virus type 1 variants in the natural course of infection. Virology 390:348–355. doi:S0042-6822(09)00335-3 [pii];10.1016/j.virol.2009.05.028 [doi].

46. Cham F, Zhang PF, Heyndrickx L, Bouma P, Zhong P, Katinger H, Robinson J, van der Groen G, Quinnan GV, Jr. 2006. Neutralization and infectivity characteristics of envelope glycoproteins from human immunodeficiency virus type 1 infected donors whose sera exhibit broadly cross-reactive neutralizing activity. Virology 347:36–51. doi:S0042-6822(05)00754-3 [pii];10.1016/j.virol.2005.11.019 [doi].

47. Dong XN, Xiao Y, Chen YH. 2001. ELNKWA-epitope specific antibodies induced by epitope-vaccine recognize ELDKWA- and other two neutralizing-resistant mutated epitopes on HIV-1 gp41. Immunol Lett 75:149–152. doi:S0165-2478(00)00298-4 [pii].

48. Dong XN, Wu Y, Chen YH. 2005. The neutralizing epitope ELDKWA on HIV-1 gp41: genetic variability and antigenicity. Immunol Lett 101:81–86. doi:S0165-2478(05)00108-2 [pii];10.1016/j.imlet.2005.04.014 [doi].

49. Gray ES, Moore PL, Bibollet-Ruche F, Li H, Decker JM, Meyers T, Shaw GM, Morris L. 2008. 4E10-resistant variants in a human immunodeficiency virus type 1 subtype C-infected individual with an anti-membrane-proximal external region-neutralizing antibody response. J Virol 82:2367–2375. doi:JVI.02161-07 [pii];10.1128/JVI.02161-07 [doi].

50. Gray ES, Madiga MC, Moore PL, Mlisana K, Abdool Karim SS, Binley JM, Shaw GM, Mascola JR, Morris L. 2009. Broad neutralization of human immunodeficiency virus type 1 mediated by plasma antibodies against the gp41 membrane proximal external region. J Virol 83:11265–11274. doi:JVI.01359-09 [pii];10.1128/JVI.01359-09 [doi].

51. Huang J, Dong X, Liu Z, Qin L, Chen YH. 2002. A predefined epitope-specific monoclonal antibody recognizes ELDEWA-epitope just presenting on gp41 of HIV-1 O clade. Immunol Lett 84:205–209. doi:S0165247802001748 [pii].

52. Huang J, Ofek G, Laub L, Louder MK, Doria-Rose NA, Longo NS, Imamichi H, Bailer RT, Chakrabarti B, Sharma SK, Alam SM, Wang T, Yang Y, Zhang B, Migueles SA, Wyatt R, Haynes BF, Kwong PD, Mascola JR, Connors M. 2012. Broad and potent neutralization of HIV-1 by a gp41-specific human antibody. Nature 491:406–412. doi:nature11544 [pii];10.1038/nature11544 [doi].

53. Li M, Salazar-Gonzalez JF, Derdeyn CA, Morris L, Williamson C, Robinson JE, Decker JM, Li Y, Salazar MG, Polonis VR, Mlisana K, Karim SA, Hong K, Greene KM, Bilska M, Zhou J, Allen S, Chomba E, Mulenga J, Vwalika C, Gao F, Zhang M, Korber BT, Hunter E, Hahn BH, Montefiori DC. 2006. Genetic and neutralization properties of subtype C human immunodeficiency virus type 1 molecular env clones from acute and early heterosexually acquired infections in Southern Africa. J Virol 80:11776–11790. doi:JVI.01730-06 [pii];10.1128/JVI.01730-06 [doi].

54. Manrique A, Rusert P, Joos B, Fischer M, Kuster H, Leemann C, Niederost B, Weber R, Stiegler G, Katinger H, Gunthard HF, Trkola A. 2007. In vivo and in vitro escape from neutralizing antibodies 2G12, 2F5, and 4E10. J Virol 81:8793–8808. doi:JVI.00598-07 [pii];10.1128/JVI.00598-07 [doi].

55. Nakowitsch S, Quendler H, Fekete H, Kunert R, Katinger H, Stiegler G. 2005. HIV-1 mutants escaping neutralization by the human antibodies 2F5, 2G12, and 4E10: in vitro experiments versus clinical studies. AIDS 19:1957–1966. doi:00002030-200511180-00003 [pii].

56. Nandi A, Lavine CL, Wang P, Lipchina I, Goepfert PA, Shaw GM, Tomaras GD, Montefiori DC, Haynes BF, Easterbrook P, Robinson JE, Sodroski JG, Yang X. 2010. Epitopes for broad and potent neutralizing antibody responses during chronic infection with human immunodeficiency virus type 1. Virology 396:339–348. doi:S0042-6822(09)00680-1 [pii];10.1016/j.virol.2009.10.044 [doi].

57. Nelson JD, Brunel FM, Jensen R, Crooks ET, Cardoso RM, Wang M, Hessell A, Wilson IA, Binley JM, Dawson PE, Burton DR, Zwick MB. 2007. An affinity-enhanced neutralizing antibody against the membrane-proximal external region of human immunodeficiency virus type 1 gp41 recognizes an epitope between those of 2F5 and 4E10. J Virol 81:4033–4043. doi:JVI.02588-06 [pii];10.1128/JVI.02588-06 [doi].

58. Pejchal R, Gach JS, Brunel FM, Cardoso RM, Stanfield RL, Dawson PE, Burton DR, Zwick MB, Wilson IA. 2009. A conformational switch in human immunodeficiency virus gp41 revealed by the structures of overlapping epitopes recognized by neutralizing antibodies. J Virol 83:8451–8462. doi:JVI.00685-09 [pii];10.1128/JVI.00685-09 [doi].

59. Shen X, Dennison SM, Liu P, Gao F, Jaeger F, Montefiori DC, Verkoczy L, Haynes BF, Alam SM, Tomaras GD. 2010. Prolonged exposure of the HIV-1 gp41 membrane proximal region with L669S substitution. Proc Natl Acad Sci U S A 107:5972–5977. doi:0912381107 [pii];10.1073/pnas.0912381107 [doi].

60. Wang Z, Liu Z, Cheng X, Chen YH. 2005. The recombinant immunogen with high-density epitopes of ELDKWA and ELDEWA induced antibodies recognizing both epitopes on HIV-1 gp41. Microbiol Immunol 49:703–709. doi:JST.JSTAGE/mandi/49.703 [pii].

61. Yang Z. 1997. PAML: a program package for phylogenetic analysis by maximum likelihood. Comput Appl Biosci 13:555–556.

62. Crooks GE, Hon G, Chandonia JM, Brenner SE. 2004. WebLogo: a sequence logo generator. Genome Res 14:1188–1190. doi:10.1101/gr.849004 [doi];14/6/1188 [pii].

63. Kyte J, Doolittle RF. 1982. A simple method for displaying the hydropathic character of a protein. J Mol Biol 157:105–132. doi:0022-2836(82)90515-0 [pii].

64. Sen J, Yan T, Wang J, Rong L, Tao L, Caffrey M. 2010. Alanine scanning mutagenesis of HIV-1 gp41 heptad repeat 1: insight into the gp120-gp41 interaction. Biochemistry 49:5057–5065. doi:10.1021/bi1005267 [doi].

65. Gonzalez N, Alvarez A, Alcami J. 2010. Broadly neutralizing antibodies and their significance for HIV-1 vaccines. Curr HIV Res 8:602–612. doi:ABS-109 [pii].

66. Koh WW, Forsman A, Hue S, van der Velden GJ, Yirrell DL, McKnight A, Weiss RA, Aasa-Chapman MM. 2010. Novel subtype C human immunodeficiency virus type 1 envelopes cloned directly from plasma: coreceptor usage and neutralization phenotypes. J Gen Virol 91:2374–2380. doi:vir.0.022228-0 [pii];10.1099/vir.0.022228-0 [doi].

67. Ringe R, Thakar M, Bhattacharya J. 2010. Variations in autologous neutralization and CD4 dependence of b12 resistant HIV-1 clade C env clones obtained at different time points from antiretroviral naive Indian patients with recent infection. Retrovirology 7:76. doi:1742-4690-7-76 [pii];10.1186/1742-4690-7-76 [doi].

68. Zwick MB. 2005. The membrane-proximal external region of HIV-1 gp41: a vaccine target worth exploring. AIDS 19:1725–1737. doi:00002030-200511040-00001 [pii].

69. Nicoll A, Killewo JZ, Mgone C. 1990. HIV and infant feeding practices: epidemiological implications for sub-Saharan African countries. AIDS 4:661–665.

70. Ross JS, Labbok MH. 2004. Modeling the effects of different infant feeding strategies on infant survival and mother-to-child transmission of HIV. Am J Public Health 94:1174–1180. doi:94/7/1174 [pii].

71. Ryder RW, Manzila T, Baende E, Kabagabo U, Behets F, Batter V, Paquot E, Binyingo E, Heyward WL. 1991. Evidence from Zaire that breast-feeding by HIV-1-seropositive mothers is not a major route for perinatal HIV-1 transmission but does decrease morbidity. AIDS 5:709–714.

72. Lanari M, Sogno VP, Natale F, Capretti MG, Serra L. 2012. Human milk, a concrete risk for infection? J Matern Fetal Neonatal Med 25 Suppl 4:75–77. doi:10.3109/14767058.2012.715009 [doi].

73. Fiscus SA, Aldrovandi GM. 2012. Virologic determinants of breast milk transmission of HIV-1. Adv Exp Med Biol 743:69–80. doi:10.1007/978-1-4614-2251-8_5 [doi].

74. Mofenson LM, McIntyre JA. 2000. Advances and research directions in the prevention of mother-to-child HIV-1 transmission. Lancet 355:2237–2244. doi:S0140-6736(00)02415-6 [pii];10.1016/S0140-6736(00)02415-6 [doi].

75. Ogundele MO, Coulter JB. 2003. HIV transmission through breastfeeding: problems and prevention. Ann Trop Paediatr 23:91–106. doi:10.1179/027249303235002161 [doi].

76. Shetty AK, Maldonado Y. 2013. Antiretroviral drugs to prevent mother-to-child transmission of HIV during breastfeeding. Curr HIV Res 11:102–125. doi:CHIVR-EPUB-20130221-3 [pii].

77. Mabuka J, Goo L, Omenda MM, Nduati R, Overbaugh J. 2013. HIV-1 maternal and infant variants show similar sensitivity to broadly neutralizing antibodies, but sensitivity varies by subtype. AIDS 27:1535–1544. doi:10.1097/QAD.0b013e32835faba5 [doi];00002030-201306190-00002 [pii].

78. Milligan C, Overbaugh J. 2014. The Role of Cell-Associated Virus in Mother-to-Child HIV Transmission. J Infect Dis 210:S631–S640. doi:jiu344 [pii];10.1093/infdis/jiu344 [doi].

79. Nakamura KJ, Cerini C, Sobrera ER, Heath L, Sinkala M, Kankasa C, Thea DM, Mullins JI, Kuhn L, Aldrovandi GM. 2013. Coverage of primary mother-to-child HIV transmission isolates by second-generation broadly neutralizing antibodies. AIDS 27:337–346. doi:10.1097/QAD.0b013e32835cadd6 [doi];00002030-201301280-00004 [pii].

80. Lorizate M, Huarte N, Saez-Cirion A, Nieva JL. 2008. Interfacial pre-transmembrane domains in viral proteins promoting membrane fusion and fission. Biochim Biophys Acta 1778:1624–1639. doi:S0005-2736(07)00477-4 [pii];10.1016/j.bbamem.2007.12.018 [doi].

81. Vishwanathan SA, Hunter E. 2008. Importance of the membrane-perturbing properties of the membrane-proximal external region of human immunodeficiency virus type 1 gp41 to viral fusion. J Virol 82:5118–5126. doi:JVI.00305-08 [pii];10.1128/JVI.00305-08 [doi].

82. Chan DI, Prenner EJ, Vogel HJ. 2006. Tryptophan- and arginine-rich antimicrobial peptides: structures and mechanisms of action. Biochim Biophys Acta 1758:1184–1202. doi:S0005-2736(06)00140-4 [pii];10.1016/j.bbamem.2006.04.006 [doi].

83. Huarte N, Lorizate M, Maeso R, Kunert R, Arranz R, Valpuesta JM, Nieva JL. 2008. The broadly neutralizing anti-human immunodeficiency virus type 1 4E10 monoclonal antibody is better adapted to membrane-bound epitope recognition and blocking than 2F5. J Virol 82:8986–8996. doi:JVI.00846-08 [pii];10.1128/JVI.00846-08 [doi].

84. Apellaniz B, Nir S, Nieva JL. 2009. Distinct mechanisms of lipid bilayer perturbation induced by peptides derived from the membrane-proximal external region of HIV-1 gp41. Biochemistry 48:5320–5331. doi:10.1021/bi900504t [doi].

85. Ionov M, Ciepluch K, Garaiova Z, Melikishvili S, Michlewska S, Balcerzak L, Glinska S, Milowska K, Gomez-Ramirez R, de la Mata FJ, Shcharbin D, Waczulikova I, Bryszewska M, Hianik T. 2015. Dendrimers complexed with HIV-1 peptides interact with liposomes and lipid monolayers. Biochim Biophys Acta 1848:907–915. doi:S0005-2736(15)00003-6 [pii];10.1016/j.bbamem.2014.12.025 [doi].

86. Kumar SB, Handelman SK, Voronkin I, Mwapasa V, Janies D, Rogerson SJ, Meshnick SR, Kwiek JJ. 2011. Different regions of HIV-1 subtype C env are associated with placental localization and in utero mother-to-child transmission. J Virol 85:7142–7152. doi:JVI.01955-10 [pii];10.1128/JVI.01955-10 [doi].

87. Ismael N, Bila D, Mariani D, Vubil A, Mabunda N, Abreu C, Jani I, Tanuri A. 2014. Genetic analysis and natural polymorphisms in HIV-1 gp41 isolates from Maputo City, Mozambique. AIDS Res Hum Retroviruses 30:603–609. doi:10.1089/AID.2013.0244 [doi].

88. Mzoughi O, Gaston F, Granados GC, Lakhdar-Ghazal F, Giralt E, Bahraoui E. 2010. Fusion intermediates of HIV-1 gp41 as targets for antibody production: design, synthesis, and HR1-HR2 complex purification and characterization of generated antibodies. ChemMedChem 5:1907–1918. doi:10.1002/cmdc.201000313 [doi].

89. Miyauchi K, Komano J, Yokomaku Y, Sugiura W, Yamamoto N, Matsuda Z. 2005. Role of the specific amino acid sequence of the membrane-spanning domain of human immunodeficiency virus type 1 in membrane fusion. J Virol 79:4720–4729. doi:79/8/4720 [pii];10.1128/JVI.79.8.4720-4729.2005 [doi].

90. Miyauchi K, Curran AR, Long Y, Kondo N, Iwamoto A, Engelman DM, Matsuda Z. 2010. The membrane-spanning domain of gp41 plays a critical role in intracellular trafficking of the HIV envelope protein. Retrovirology 7:95. doi:1742-4690-7-95 [pii];10.1186/1742-4690-7-95 [doi].

91. Los Alamos National Laboratory. HIV molecular immunology database. Mar 31, 2015. http://www.hiv.lanl.gov/content/immunology/tables/ab_summary.html

## REFERENCES FOR SUPPLEMENTAL FIGURES

1. Bunnik EM, van Gils MJ, Lobbrecht MS, Pisas L, van Nuenen AC, Schuitemaker H. 2009. Changing sensitivity to broadly neutralizing antibodies b12, 2G12, 2F5, and 4E10 of primary subtype B human immunodeficiency virus type 1 variants in the natural course of infection. Virology 390:348–355. doi:S0042-6822(09)00335-3 [pii];10.1016/j.virol.2009.05.028 [doi].

2. Dong XN, Xiao Y, Chen YH. 2001. ELNKWA-epitope specific antibodies induced by epitope-vaccine recognize ELDKWA- and other two neutralizing-resistant mutated epitopes on HIV-1 gp41. Immunol Lett 75:149–152. doi:S0165-2478(00)00298-4 [pii].

3. Dong XN, Wu Y, Chen YH. 2005. The neutralizing epitope ELDKWA on HIV-1 gp41: genetic variability and antigenicity. Immunol Lett 101:81–86. doi:S0165-2478(05)00108-2 [pii];10.1016/j.imlet.2005.04.014 [doi].

4. Gray ES, Moore PL, Bibollet-Ruche F, Li H, Decker JM, Meyers T, Shaw GM, Morris L. 2008. 4E10-resistant variants in a human immunodeficiency virus type 1 subtype C-infected individual with an anti-membrane-proximal external region-neutralizing antibody response. J Virol 82:2367–2375. doi:JVI.02161-07 [pii];10.1128/JVI.02161-07 [doi].

5. Gray ES, Madiga MC, Moore PL, Mlisana K, Abdool Karim SS, Binley JM, Shaw GM, Mascola JR, Morris L. 2009. Broad neutralization of human immunodeficiency virus type 1 mediated by plasma antibodies against the gp41 membrane proximal external region. J Virol 83:11265–11274. doi:JVI.01359-09 [pii];10.1128/JVI.01359-09 [doi].

6. Huang J, Dong X, Liu Z, Qin L, Chen YH. 2002. A predefined epitope-specific monoclonal antibody recognizes ELDEWA-epitope just presenting on gp41 of HIV-1 O clade. Immunol Lett 84:205–209. doi:S0165247802001748 [pii].

7. Huang J, Ofek G, Laub L, Louder MK, Doria-Rose NA, Longo NS, Imamichi H, Bailer RT, Chakrabarti B, Sharma SK, Alam SM, Wang T, Yang Y, Zhang B, Migueles SA, Wyatt R, Haynes BF, Kwong PD, Mascola JR, Connors M. 2012. Broad and potent neutralization of HIV-1 by a gp41-specific human antibody. Nature 491:406–412. doi:nature11544 [pii];10.1038/nature11544 [doi].

8. Manrique A, Rusert P, Joos B, Fischer M, Kuster H, Leemann C, Niederost B, Weber R, Stiegler G, Katinger H, Gunthard HF, Trkola A. 2007. In vivo and in vitro escape from neutralizing antibodies 2G12, 2F5, and 4E10. J Virol 81:8793–8808. doi:JVI.00598-07 [pii];10.1128/JVI.00598-07 [doi].

9. Nakamura KJ, Gach JS, Jones L, Semrau K, Walter J, Bibollet-Ruche F, Decker JM, Heath L, Decker WD, Sinkala M, Kankasa C, Thea D, Mullins J, Kuhn L, Zwick MB, Aldrovandi GM. 2010. 4E10-resistant HIV-1 isolated from four subjects with rare membrane-proximal external region polymorphisms. PLoS One 5:e9786. doi:10.1371/journal.pone.0009786 [doi].

10. Nakowitsch S, Quendler H, Fekete H, Kunert R, Katinger H, Stiegler G. 2005. HIV-1 mutants escaping neutralization by the human antibodies 2F5, 2G12, and 4E10: in vitro experiments versus clinical studies. AIDS 19:1957–1966. doi:00002030-200511180-00003 [pii].

11. Nandi A, Lavine CL, Wang P, Lipchina I, Goepfert PA, Shaw GM, Tomaras GD, Montefiori DC, Haynes BF, Easterbrook P, Robinson JE, Sodroski JG, Yang X. 2010. Epitopes for broad and potent neutralizing antibody responses during chronic infection with human immunodeficiency virus type 1. Virology 396:339–348. doi:S0042-6822(09)00680-1 [pii];10.1016/j.virol.2009.10.044 [doi].

12. Nelson JD, Brunel FM, Jensen R, Crooks ET, Cardoso RM, Wang M, Hessell A, Wilson IA, Binley JM, Dawson PE, Burton DR, Zwick MB. 2007. An affinity-enhanced neutralizing antibody against the membrane-proximal external region of human immunodeficiency virus type 1 gp41 recognizes an epitope between those of 2F5 and 4E10. J Virol 81:4033–4043. doi:JVI.02588-06 [pii];10.1128/JVI.02588-06 [doi].

13. Pejchal R, Gach JS, Brunel FM, Cardoso RM, Stanfield RL, Dawson PE, Burton DR, Zwick MB, Wilson IA. 2009. A conformational switch in human immunodeficiency virus gp41 revealed by the structures of overlapping epitopes recognized by neutralizing antibodies. J Virol 83:8451–8462. doi:JVI.00685-09 [pii];10.1128/JVI.00685-09 [doi].

14. Wang Z, Liu Z, Cheng X, Chen YH. 2005. The recombinant immunogen with high-density epitopes of ELDKWA and ELDEWA induced antibodies recognizing both epitopes on HIV-1 gp41. Microbiol Immunol 49:703–709. doi:JST.JSTAGE/mandi/49.703 [pii].

15. Zwick MB, Jensen R, Church S, Wang M, Stiegler G, Kunert R, Katinger H, Burton DR. 2005. Anti-human immunodeficiency virus type 1 (HIV-1) antibodies 2F5 and 4E10 require surprisingly few crucial residues in the membrane-proximal external region of glycoprotein gp41 to neutralize HIV-1. J Virol 79:1252–1261. doi:79/2/1252 [pii];10.1128/JVI.79.2.1252-1261.2005 [doi].

16. Blish CA, Nguyen MA, Overbaugh J. 2008. Enhancing exposure of HIV-1 neutralization epitopes through mutations in gp41. PLoS Med 5:e9. doi:07-PLME-RA-0379 [pii];10.1371/journal.pmed.0050009 [doi].

17. Cham F, Zhang PF, Heyndrickx L, Bouma P, Zhong P, Katinger H, Robinson J, van der Groen G, Quinnan GV, Jr. 2006. Neutralization and infectivity characteristics of envelope glycoproteins from human immunodeficiency virus type 1 infected donors whose sera exhibit broadly cross-reactive neutralizing activity. Virology 347:36–51. doi:S0042-6822(05)00754-3 [pii];10.1016/j.virol.2005.11.019 [doi].

18. Li M, Salazar-Gonzalez JF, Derdeyn CA, Morris L, Williamson C, Robinson JE, Decker JM, Li Y, Salazar MG, Polonis VR, Mlisana K, Karim SA, Hong K, Greene KM, Bilska M, Zhou J, Allen S, Chomba E, Mulenga J, Vwalika C, Gao F, Zhang M, Korber BT, Hunter E, Hahn BH, Montefiori DC. 2006. Genetic and neutralization properties of subtype C human immunodeficiency virus type 1 molecular env clones from acute and early heterosexually acquired infections in Southern Africa. J Virol 80:11776–11790. doi:JVI.01730-06 [pii];10.1128/JVI.01730-06 [doi].

19. Shen X, Dennison SM, Liu P, Gao F, Jaeger F, Montefiori DC, Verkoczy L, Haynes BF, Alam SM, Tomaras GD. 2010. Prolonged exposure of the HIV-1 gp41 membrane proximal region with L669S substitution. Proc Natl Acad Sci U S A 107:5972–5977. doi:0912381107 [pii];10.1073/pnas.0912381107 [doi].

20. Los Alamos National Laboratory. HIV molecular immunology database. 3-31-2015. http://www.hiv.lanl.gov/content/immunology/tables/ab_summary.html

21. Ratner L, Haseltine W, Patarca R, Livak KJ, Starcich B, Josephs SF, Doran ER, Rafalski JA, Whitehorn EA, Baumeister K,. 1985. Complete nucleotide sequence of the AIDS virus, HTLV-III. Nature 313:277–284.

22. Perez-Alvarez L, Carmona R, Ocampo A, Asorey A, Miralles C, Perez de CS, Pinilla M, Contreras G, Taboada JA, Najera R. 2006. Long-term monitoring of genotypic and phenotypic resistance to T20 in treated patients infected with HIV-1. J Med Virol 78:141–147. doi: 10.1002/jmv.20520 [doi].

23. Si-Mohamed A, Piketty C, Tisserand P, LeGoff J, Weiss L, Charpentier C, Kazatchkine MD, Belec L. 2007. Increased polymorphism in the HR-1 gp41 env gene encoding the enfuvirtide (T-20) target in HIV-1 variants harboring multiple antiretroviral drug resistance mutations in the pol gene. J Acquir Immune Defic Syndr 44:1–5. doi:10.1097/01.qai.0000243118.59906.f4 [doi].

24. Teixeira C, de Sa-Filho D, Alkmim W, Janini LM, Diaz RS, Komninakis S. 2010. Short communication: high polymorphism rates in the HR1 and HR2 gp41 and presence of primary resistance-related mutations in HIV type 1 circulating in Brazil: possible impact on enfuvirtide efficacy. AIDS Res Hum Retroviruses 26:307–311. doi:10.1089/aid.2008.0297 [doi].

25. Long Y, Meng F, Kondo N, Iwamoto A, Matsuda Z. 2011. Conserved arginine residue in the membrane-spanning domain of HIV-1 gp41 is required for efficient membrane fusion. Protein Cell 2:369–376. doi:10.1007/s13238-011-1051-0 [doi].

26. Miyauchi K, Komano J, Yokomaku Y, Sugiura W, Yamamoto N, Matsuda Z. 2005. Role of the specific amino acid sequence of the membrane-spanning domain of human immunodeficiency virus type 1 in membrane fusion. J Virol 79:4720–4729. doi:79/8/4720 [pii];10.1128/JVI.79.8.4720-4729.2005 [doi].

27. Miyauchi K, Curran AR, Long Y, Kondo N, Iwamoto A, Engelman DM, Matsuda Z. 2010. The membrane-spanning domain of gp41 plays a critical role in intracellular trafficking of the HIV envelope protein. Retrovirology 7:95. doi:1742-4690-7-95 [pii];10.1186/1742-4690-7-95 [doi].

